# Light-dependent changes in the higher-order DNA structure of the cyanobacterium *Synechocystis* sp. PCC6803

**DOI:** 10.64898/2026.04.09.717459

**Authors:** Ryo Kariyazono, Hideyuki Tanabe, Takashi Osanai

**Author notes:** Corresponding author; Ryo Kariyazono email. Corresponding author; Takashi Osanai email.

## Abstract

Chromosome spatial organization plays critical roles in transcriptional regulation and DNA protection. In cyanobacteria—photosynthetic bacteria that experience dramatic fluctuations in light intensity—chromosome reorganization could facilitate rapid transcriptional reprogramming and protect DNA from photodamage. However, chromosome organization in these polyploid organisms has remained technically challenging to observe, leaving light-dependent responses unexplored.

Here, we show that higher-order chromosome organization in *Synechocystis* sp. PCC 6803 is associated with light intensity, revealing a previously unrecognized light-dependent adaptation in cyanobacteria. We established fluorescence in situ hybridization (FISH) methods for this model cyanobacterium carrying multi-copy genomes, together with a computational pipeline to assign paired FISH signals to individual genome copies. The slope relating genomic and spatial distance was steeper under standard conditions (β = 0.972 nm/kbp, R² = 0.12) than under high-light conditions (β = 0.450 nm/kbp, R² = 0.02), indicating that local chromosome organization is substantially disrupted by elevated light intensity. The spatial distribution of the multiple genome copies also differed between conditions, independently supporting condition-dependent chromosome reorganization. Hi-C analysis corroborated these findings, revealing reduced chromosomal interactions within the 10–100 kbp range under high-light conditions. Together, these results demonstrate that light intensity is a previously unrecognized determinant of higher-order chromosome organization in a photosynthetic bacterium.

## INTRODUCTION

Higher-order chromosome organization plays critical roles in transcriptional regulation and DNA protection across both eukaryotes and prokaryotes (Hołówka & Zakrzewska-Czerwińska, 2020; Walker *et al*, 2025). Cyanobacteria—oxygenic photosynthetic bacteria—experience dramatic fluctuations in light intensity in natural environments, and must rapidly reprogram transcription and protect their DNA from photodamage in response to changing light conditions. These responses encompass a broad repertoire of molecular mechanisms, including metabolic reprogramming during diurnal cycles(Welkie *et al*, 2019), chromatic acclimation to shifts in light quality (Sanfilippo *et al*, 2019), and photoprotection under high-light stress (Muzzopappa & Kirilovsky, 2020). In particular, light intensity drives large-scale transcriptional reprogramming(Muramatsu & Hihara, 2012) and induces DNA damage that requires active repair mechanisms(Singh *et al*, 2023)—two processes in which higher-order chromosome organization is known to play a central role (Walker *et al*, 2025). Consistent with this idea, recent 3C and Hi-C studies have shown that cyanobacterial chromosome organization responds to environmental changes including microoxic conditions and iron depletion(Kariyazono & Osanai, 2024; Liu *et al*, 2024), raising the possibility that light intensity may similarly modulate chromosome structure. However, this has remained unexplored.

Polyploidy is a common feature of cyanobacteria. Cyanobacteria inhabit across marine, freshwater, and terrestrial environments, where marine cyanobacteria tend to be monoploid or oligoploid (two to four copies) and freshwater species are predominantly oligoploid to polyploid (Griese *et al*, 2011; Weissenbach *et al*, 2024). Furthermore, the genome copy number (GCN) varies in response to environmental conditions. Notably, the well-studied model freshwater cyanobacterium *Synechocystis* sp. PCC 6803 is polyploid under phosphorus-replete conditions but becomes monoploid under phosphorus-depleted conditions (Zerulla *et al*, 2016).

Due to cyanobacterial polyploidy, Hi-C is insufficient to characterize the higher-order DNA structure comprehensively. This limitation arises because polyploidy creates inter-copy locus interactions that Hi-C cannot distinguish. Furthermore, Hi-C analysis averages out inter-copy structural heterogeneity and copy number fluctuations.

Fluorescence in situ hybridization (FISH) overcomes these limitations as it offers direct visualization of higher-order DNA shape at the single-cell level. FISH enables visualization of individual copies of genomic regions, allowing detection of structural heterogeneity among genomic copies averaged out in Hi-C analysis. This approach provides complementary information to Hi-C data, particularly helpful for polyploid organisms.

Another approach to visualizing genomic loci involves lacO tandem repeats and GFP-fused lacI repressor systems. This method enables live imaging and provides an easy way to count genome ploidy and the spatial organization of multi-copy chromosomes in cyanobacteria (Jain *et al*, 2012). However, this system does not visualize endogenous genomic loci as it requires artificial insertion of a tandem lacO repeat into the genomic region of interest. Therefore, to observe changes in the higher-order DNA structure, FISH is preferable.

Although researchers have applied FISH methods for localizing genomic regions to some polyploid bacteria, including the rod-shaped cyanobacterium *Synechococcus elongatus* (Takacs *et al*, 2022; Watanabe *et al*, 2018), their application to coccoid cyanobacteria such as *Synechocystis* sp. PCC 6803 remains challenging. The spherical cell morphology and higher degree of polyploidy of *Synechocystis* make it more difficult to resolve and assign individual genomic signals compared with rod-shaped species. Moreover, unlike well-characterized model bacteria such as *Escherichia coli*—where chromosome organization can be anchored to landmarks such as the predictable GCN of one to two copies per cell and the polar localization of the replication origin (Niki *et al*, 2000)—cyanobacteria lack equivalent validation landmarks, as their replication and chromosome segregation are decoupled (Watanabe *et al*, 2012). Nevertheless, investigating chromosome organization in *Synechocystis* is valuable precisely because of this morphological contrast: features shared between rod-shaped and coccoid cyanobacteria are more likely to represent universal principles of cyanobacterial chromosome organization. In this study, we established a FISH method to localize genomic loci within *Synechocystis* sp. PCC 6803. We validated the method using cells under conditions of varying ploidy (polyploid to oligoploid) and several pairs of probes with different genomic distances. Subsequently, we confirmed that the ploidy calculated from cell counts and qPCR matched the mean number of FISH signals per cell, and that genomic and spatial distances across multiple loci were positively correlated.

Here, we report that higher-order chromosome organization in *Synechocystis* sp. PCC 6803 changes in response to high-light conditions. Using two-color FISH combined with Hi-C analysis, we demonstrate that the genomic–spatial distance relationship is substantially weakened under high-light stress, and that the spatial distribution of genome copies shifts toward a more random arrangement. These findings reveal a previously unrecognized association between light intensity and higher-order chromosome organization in cyanobacteria, and establish an integrative single-cell and population-level framework for studying chromosome organization in polyploid bacteria.

## RESULTS

### Overview of FISH analysis

We performed two-color FISH using Spectrum™ Green and Spectrum™ Orange probes on *Synechocystis* grown under standard conditions. The green probe was centered at genome position 1,702,186 bp (NC_000911.1), while the orange probes were centered at positions 25.3 kbp upstream, or 53.7, 73.6, or 123.7 kbp downstream (Figure 1A). For each experiment, the cells were hybridized with the green probe mixed with one of the four orange probes. In all probe combinations tested, we observed multiple green and orange signals per cell (Figure 1B), reflecting the multiple genome copies in *Synechocystis* cells.

**Figure 1.**
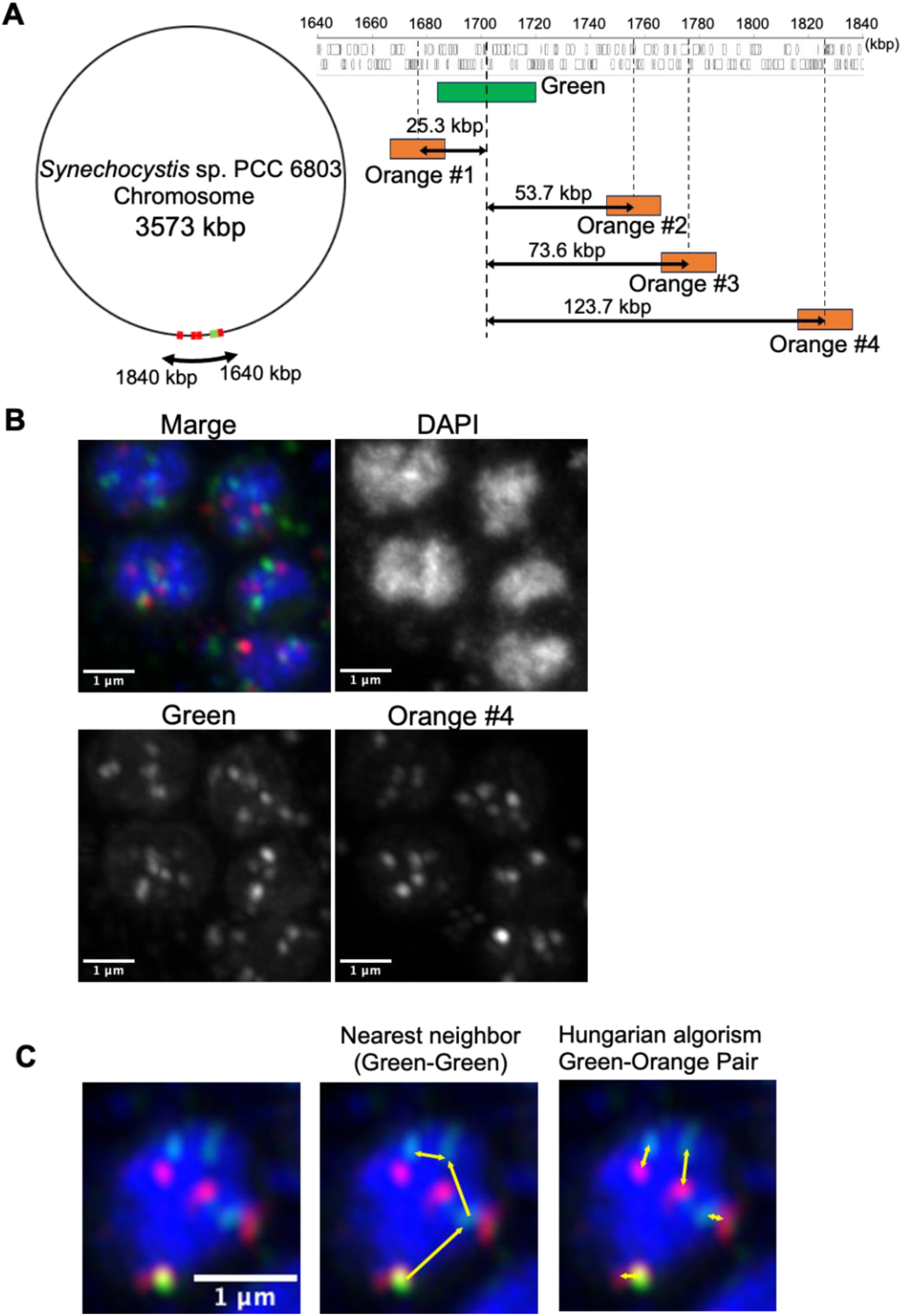
Overview of FISH experiments. **(A)** Genomic position of probes used in this study. The left image shows the position of probes within the whole chromosome of the sp. PCC 6803, and the right is a magnified image. **(B)** Snapshot of the FISH experiment. The scale bar: 1µm. **(C)** Schematic image of the nearest-neighbor distance between the same color signals (middle) and the green-orange signal pair assigned by the Hungarian algorithm (right). The scale bar: 1µm.

Each FISH experiment was independently repeated twice using biological replicates. Data from both replicates were pooled for statistical analysis.

Two complementary spatial distance measurements were conducted to characterize chromosome organization. First, to assess the spatial distribution of genome copies, we measured nearest-neighbor distances between signals of the same color—green–green and orange–orange distances (Figure 1C middle). These same-color distances reflect how chromosome copies are spatially arranged within cells, as each signal of a given color represents the same genomic locus on different copies. Second, to examine spatial distances between different genomic loci on the same chromosome, we paired green and orange signals using the Hungarian algorithm, which identifies the pairing configuration that minimizes the total sum of inter-probe distances (Figure 1C right). This pairing algorithm was validated through the following analyses.

### FISH signal count reflects the cell ploidy

To validate FISH for genome visualization, we compared FISH signal counts and copy number calculated from qPCR. As *Synechocystis* reduces its ploidy under phosphate-depleted conditions (Zeller *et al*, 2016), we counted FISH signals in cells from standard and phosphate-depleted cultures, representing high- and low-ploidy levels, respectively.

Under standard conditions, cells displayed 5∼6 FISH signals per cell with every probe, whereas qPCR estimated ∼6 copies/cell on average (Table 1). In 89.1% of observed cells, the difference in signal counts between green and orange signals was two or less. Under phosphate-depleted conditions, FISH signals were reduced to ∼2.5 per cell, consistent with qPCR estimates of fewer than 3 copies/cell (Table 1). Notably, the proportion of FISH-positive cells was markedly reduced under phosphate-depleted conditions. Results were consistent across two independent biological replicates (Table 1).

**Table 1.**
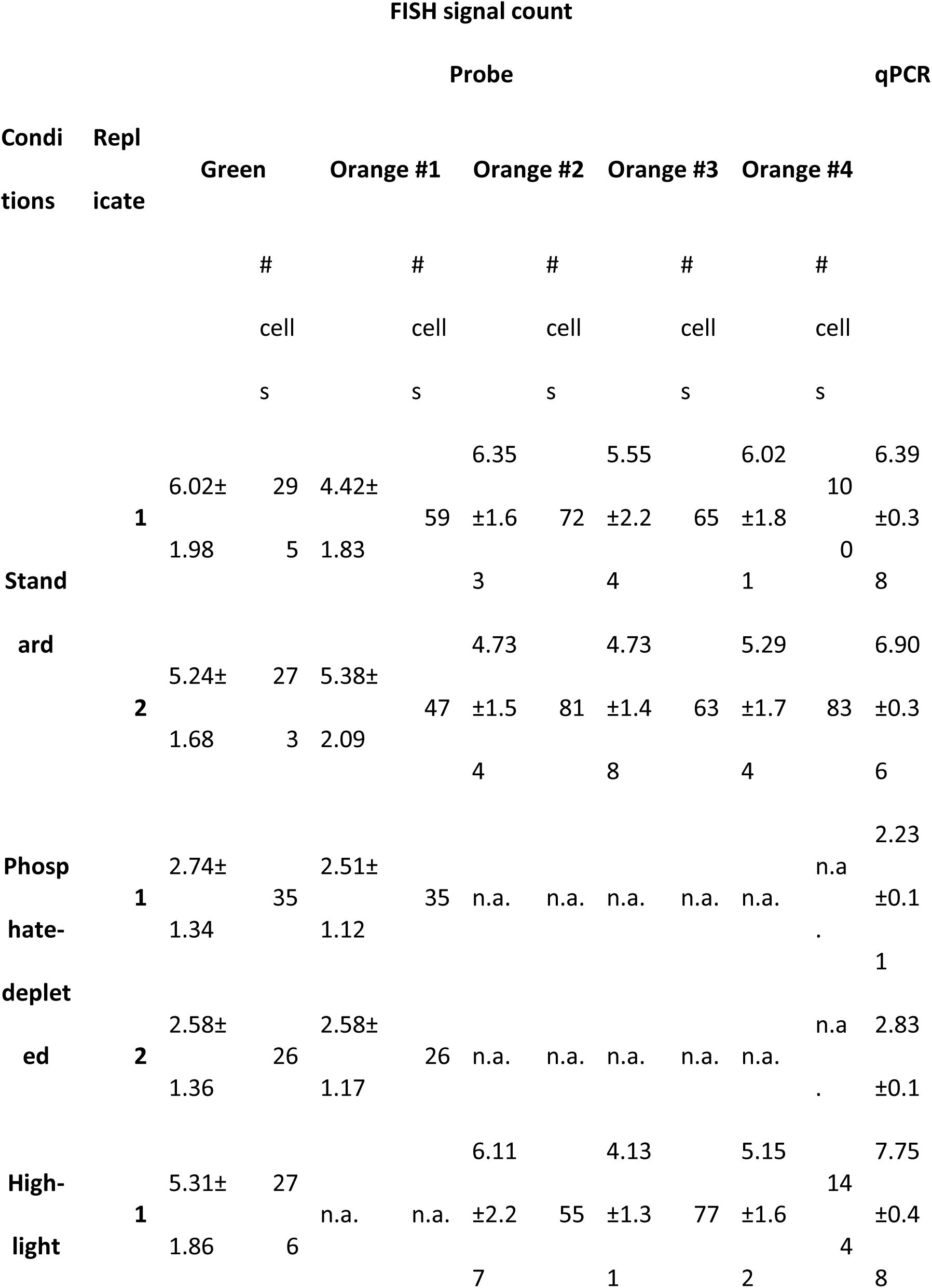
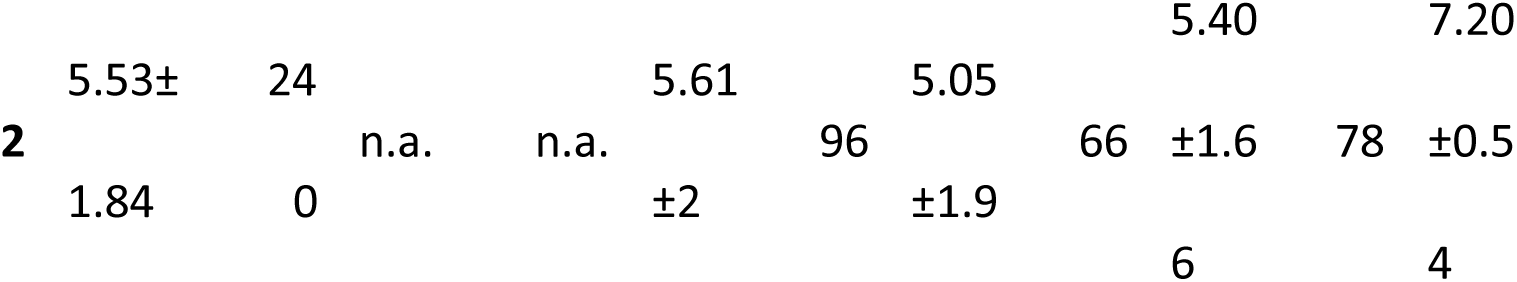
FISH signal count and qPCR-based estimated ploidy.

### Spatial distribution of genome copies

To characterize the spatial organization of genome copies within cells, we analyzed nearest-neighbor distances between signals of the same color (green–green and orange–orange), representing the same genomic locus on different chromosome copies (Figure 1C middle). Log–log linear regression of median nearest-neighbor distance versus signal number yielded slopes of −0.16 to -0.22 for green–green distances, and −0.23 to −0.28 for orange probes (pooled replicates, Table 2).

**Table 2.**
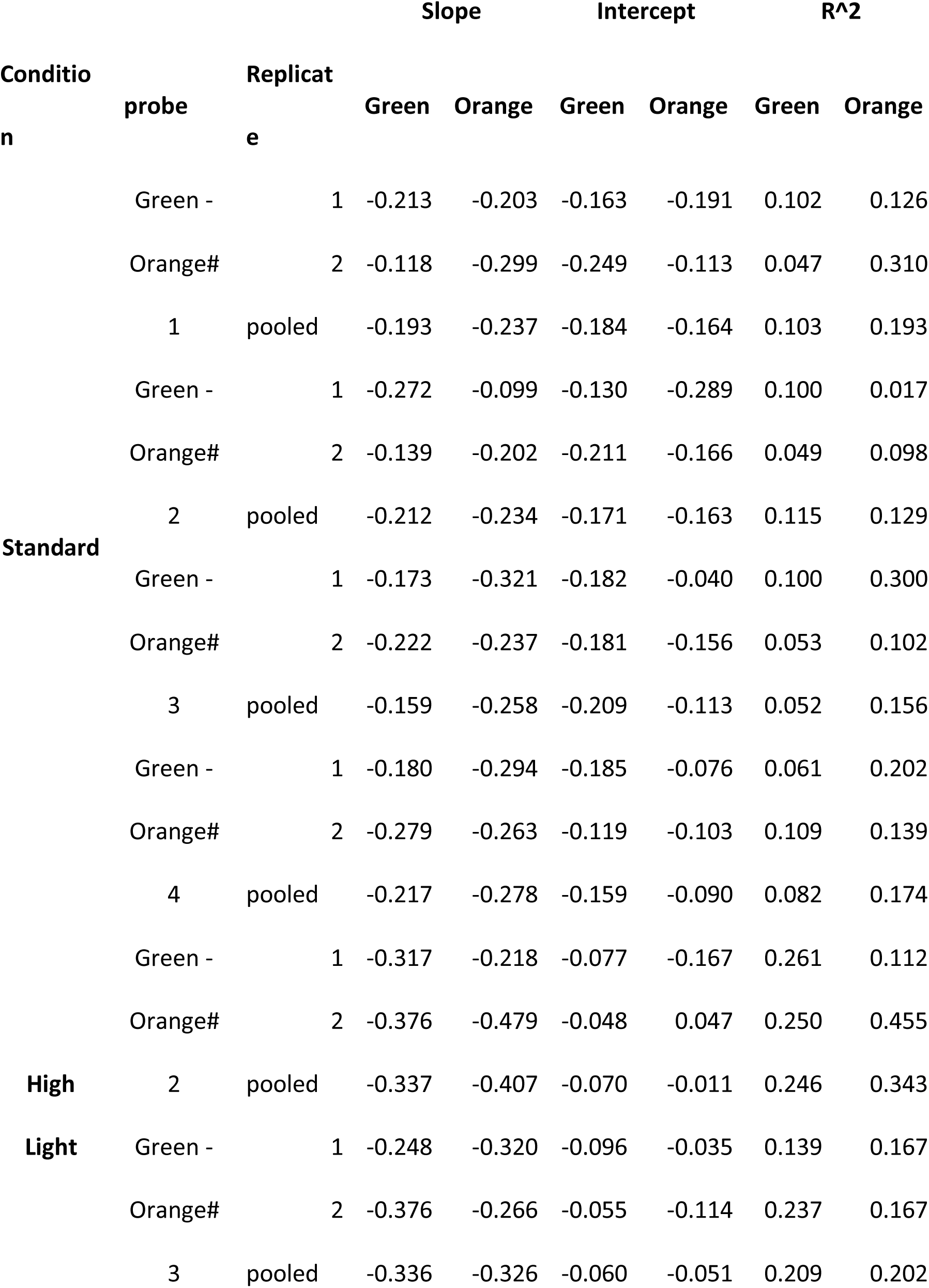
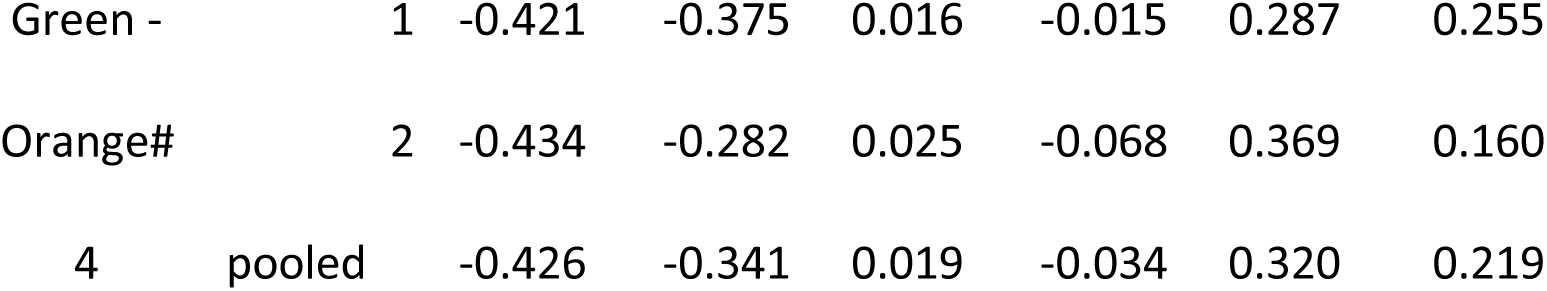
Slope of nearest-neighbor distance vs number of signals.

| Conditio<br>n | probe | Replicat<br>e | Slope |  | Intercept |  | R<sup>2</sup> |  |
| --- | --- | --- | --- | --- | --- | --- | --- | --- |
|  |  |  | Green | Orange | Green | Orange | Green | Orange |
| Standard | Green - | 1 | -0.213 | -0.203 | -0.163 | -0.191 | 0.102 | 0.126 |
|  | Orange# | 2 | -0.118 | -0.299 | -0.249 | -0.113 | 0.047 | 0.310 |
|  | 1 | pooled | -0.193 | -0.237 | -0.184 | -0.164 | 0.103 | 0.193 |
|  | Green - | 1 | -0.272 | -0.099 | -0.130 | -0.289 | 0.100 | 0.017 |
|  | Orange# | 2 | -0.139 | -0.202 | -0.211 | -0.166 | 0.049 | 0.098 |
|  | 2 | pooled | -0.212 | -0.234 | -0.171 | -0.163 | 0.115 | 0.129 |
|  | Green - | 1 | -0.173 | -0.321 | -0.182 | -0.040 | 0.100 | 0.300 |
|  | Orange# | 2 | -0.222 | -0.237 | -0.181 | -0.156 | 0.053 | 0.102 |
|  | 3 | pooled | -0.159 | -0.258 | -0.209 | -0.113 | 0.052 | 0.156 |
|  | Green - | 1 | -0.180 | -0.294 | -0.185 | -0.076 | 0.061 | 0.202 |
|  | Orange# | 2 | -0.279 | -0.263 | -0.119 | -0.103 | 0.109 | 0.139 |
|  | 4 | pooled | -0.217 | -0.278 | -0.159 | -0.090 | 0.082 | 0.174 |
| High | Green - | 1 | -0.317 | -0.218 | -0.077 | -0.167 | 0.261 | 0.112 |
|  | Orange# | 2 | -0.376 | -0.479 | -0.048 | 0.047 | 0.250 | 0.455 |
| Light | 2 | pooled | -0.337 | -0.407 | -0.070 | -0.011 | 0.246 | 0.343 |
|  | Green - | 1 | -0.248 | -0.320 | -0.096 | -0.035 | 0.139 | 0.167 |
|  | Orange# | 2 | -0.376 | -0.266 | -0.055 | -0.114 | 0.237 | 0.167 |
|  | 3 | pooled | -0.336 | -0.326 | -0.060 | -0.051 | 0.209 | 0.202 |

|  |  |  |  |  |  |  |  |
| --- | --- | --- | --- | --- | --- | --- | --- |
| Green - | 1 | -0.421 | -0.375 | 0.016 | -0.015 | 0.287 | 0.255 |
| Orange# | 2 | -0.434 | -0.282 | 0.025 | -0.068 | 0.369 | 0.160 |
| 4 | pooled | -0.426 | -0.341 | 0.019 | -0.034 | 0.320 | 0.219 |

### Spatial distance of two FISH signals corresponds to genomic distance

We next examined whether the spatial (3D) separation of genomic loci corresponds to their linear (1D) genomic distance. We mixed the common green-labeled probe separately with four orange-labeled probes positioned 25.3, 53.7, 73.6, and 123.7 kbp away and tested in the hybridization (Figure 1A). The linear regression analysis shows a significant positive correlation between genomic and spatial distances (slope β = 0.972 nm/kbp, p < 1 × 10^−15^, R² = 0.116; Figure 2A).

**Figure 2.**
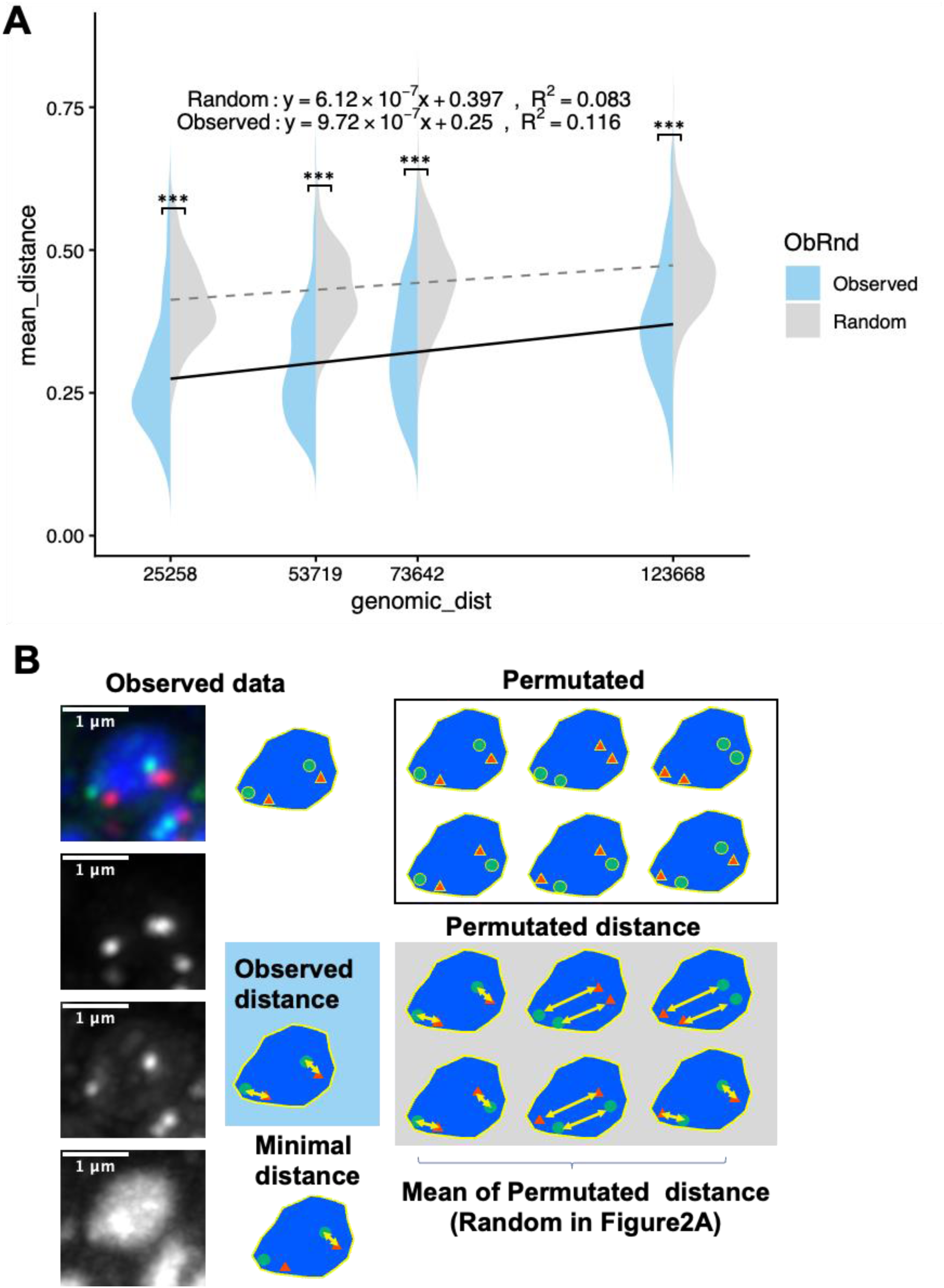
The relationship between spatial distance and genomic distance. **(A)** The relationship between spatial distance and genomic distance of the green-orange signal pair (left half violin plot for observed distance and right half violin plot for permuted distance). The solid line indicates the regression line for the observed distance, and the dashed line indicates the regression line for the permuted distance. Triple asterisks indicate p < 2.2 × 10^−16^ by paired Wilcoxon signed-rank test. **(B)** Schematic image of the calculation of the permuted pair distance. Most left is the original FISH image, and the top of middle column is an extract of green signals (green circles) and orange signals (orange triangles). The arrow in center shaded in blue is the pair distances applied with the Hungarian algorithm (the observed distance). The arrow in the bottom of middle column indicates minimal distance between green and orange signals in the cell. The right upper is all permutations of two green signals and two orange signals. The bottom right is the distances applied with the Hungarian algorithm for all permutations (the random distance).

To validate the pairing algorithm, we performed a permutation test in which the spatial coordinates were fixed but color labels (green vs. orange) were randomly reassigned to these coordinates (Figure 2B). The observed mean spatial distance of pairs per cell is significantly shorter than that from randomized data (Figure 2A). The mean spatial distance of pairs per cell from a permutation test shows a significant correlation with the genomic distances of the probes, but with a smaller slope and R^2^ than that from the observed data (slope β = 0.612 nm/kbp, p < 1 × 10⁻^11^, R^2^ = 0.083, Figure 2B). This indicates that our pairing algorithm reflects biological chromosome organization rather than geometric artifacts. To assess the robustness of these findings to potential mispairing by the Hungarian algorithm, we repeated the analysis using the minimum Green-Orange distance per cell as a pairing-independent metric (Figure 2B bottom). This alternative analysis using the minimum Green-Orange distance per cell also showed a statistically significant positive correlation between genomic and spatial distance (slope = 0.490 nm/kbp, p < 1 × 10^−9^, R² = 0.064; Supplementary Figure S1), supporting the robustness of this finding to the choice of pairing metric.

### Genome copy number does not confound spatial distance

To examine whether the genomic-spatial distance relationship varied with GCN, we fitted interaction models using the number of probe pairs per cell—an observational proxy for GCN—and genomic distance as predictors, with both mean and minimum Green-Orange distances as response variables. The interaction term was not significant in either model (mean distance: p = 0.103; minimum distance: p = 0.566, Supplementary Table S1). Notably, the number of probe pairs showed opposite effects on the two distance metrics: a positive effect on mean distance (β = +1.08×10⁻², p = 1.78×10⁻⁵) and a negative effect on minimum distance (β = −1.10×10⁻², p = 2.35×10⁻¹⁰) (Supplementary Figure S2).

### Chromosome organization under high-light conditions

To examine whether chromosome spatial organization changes under different growth conditions, we performed FISH analysis of cells cultured under high-light conditions (80 minutes of radiation by 300 μmol photons m⁻² s⁻¹). Quantitative PCR analysis estimated the GCN under high-light conditions to be ∼7 per cell. Both replicates slightly increased their ploidy compared to those under standard conditions (∼6 copies; Table 1). For high-light conditions, we hybridized cells with the green probe and one of three orange probes (53.7, 73.6, or 123.7 kbp downstream); the 25.3 kbp upstream probe was not included in high-light experiments.

For genome copy distribution (same-color nearest-neighbor distances), green–green distances showed a slope of −0.34 to -0.43 under high-light and −0.25 under standard conditions (Figure 3A). Orange probes showed similar shifts (from −0.23 to −0.28 in standard conditions to −0.33 to -0.41 for both in high-light conditions; Figure 3A).

**Figure 3.**
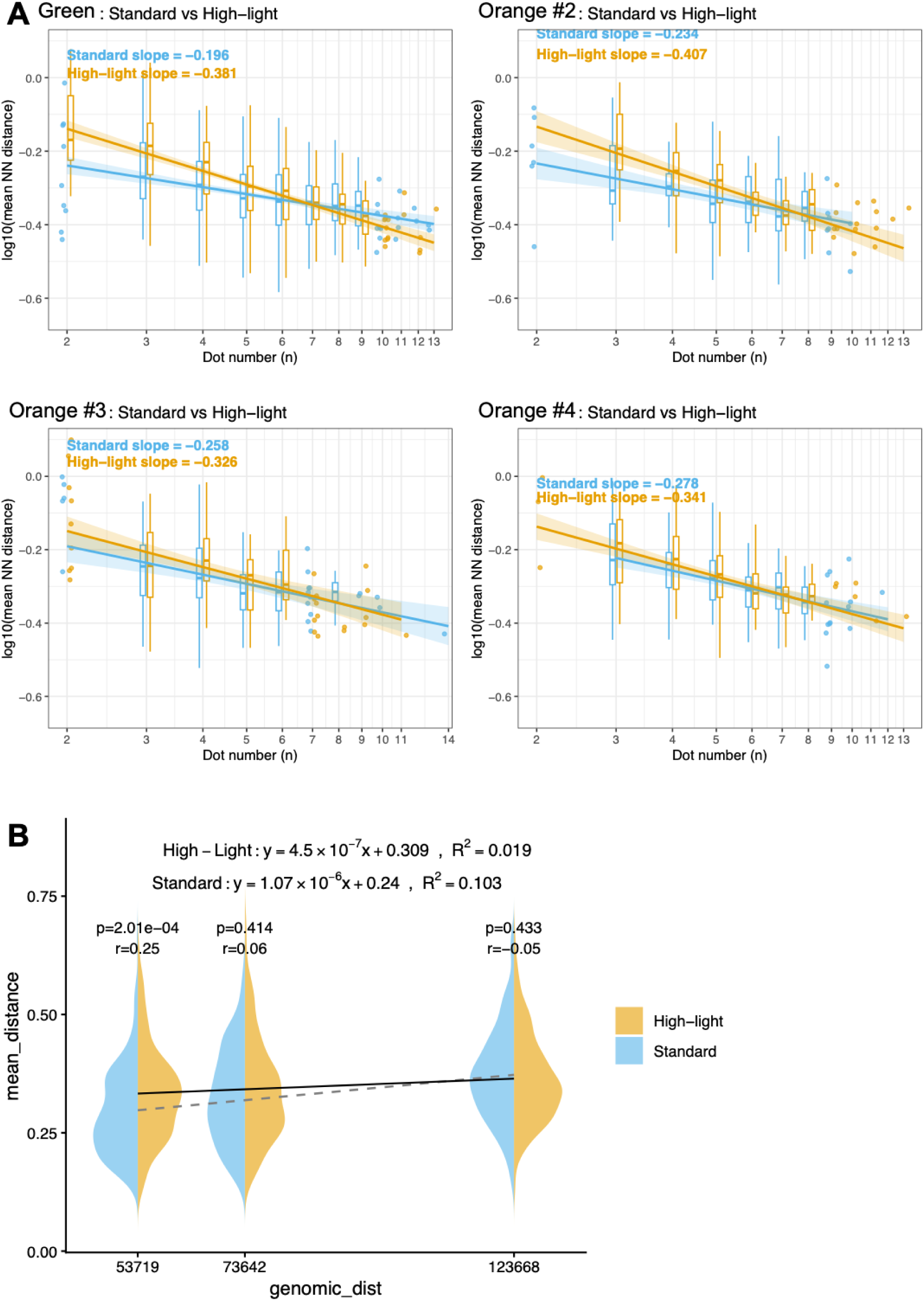
Chromosome reorganization upon the high-light exposure **(A)** Log–log plots of mean nearest-neighbor distance versus signal count per cell, shown separately for green–green (upper left) and orange–orange (orange probes #2–#4) same-color signal pairs. Each panel shows cells under standard (blue) and high-light (orange) conditions. The x-axis represents the number of same-color signals per cell (a proxy for genome copy number), and the y-axis represents the mean nearest-neighbor distance. Groups with ≥10 cells are shown as box plots; groups with <10 cells are shown as jittered points. Regression lines with 95% confidence intervals are shown for each condition. **(B)** The relationship between spatial distance and genomic distance of the green-orange signal pair (red violin plot for distance in the standard conditions and light gray violin plot for distance in the high-light condition). The solid line indicates the regression line in the high-light condition, and the dashed line indicates the regression line in the standard condition. Mann–Whitney p-values and rank-biserial correlation coefficients (r) are indicated on the violin plots.

Linear regression analysis using the three probe pairs common to both conditions (53.7, 73.6, and 123.7 kbp) revealed a significant positive correlation under standard conditions (β = 1.069 nm/kbp, R² = 0.103), which was substantially reduced under high-light conditions (β = 0.450 nm/kbp, R² = 0.019, Figure 3B). Mann−Whitney tests revealed a significant difference with small effect size in mean distances at 53.7 kbp (p = 2.0 × 10⁻^4^, r=0.25), and no significant difference at 73.6 kbp and 123.7 kbp (p = 0.414 and 0.433, r = 0.058 and −0.045, respectively) (Figure 3B). Analysis using minimum Green-Orange distances per cell yielded qualitatively consistent results: a reduced slope under high-light conditions (standard: β = 0.604 nm/kbp; high-light: β = 0.408 nm/kbp) and a significant difference at 53.7 kbp (Mann-Whitney test, p = 0.007; Supplementary Figure S3). To examine whether this altered relationship was attributable to the slightly elevated GCN under high-light (∼7 vs. ∼6 copies per cell), we fitted interaction models using the number of probe pairs per cell—an observational proxy for GCN—and genomic distance as predictors. Interaction models confirmed that this relationship was not modulated by the number of probe pairs under high-light conditions (mean distance: p = 0.156; minimum distance: p = 0.558), and the opposing effects of probe pair number on mean and minimum distances were also replicated (Supplementary Figure S2 and Supplementary Table S1).

### Hi-C analysis supports altered chromosome organization under high-light conditions

To assess the microscopic observation with a different method, we performed Hi-C analysis on *Synechocystis* under the standard condition and the high-light condition. Note that the light quantity at the high-light condition in this experiment is 150µmol photons m-^2^ s^-1^. When we plotted genomic distance vs contact frequency of cells under the standard condition, we found the shoulder at around 10 kbp to 100kbp, where the slope is shallower than in other regions (log–log slope in the 10–100 kbp range: −0.639 ± 0.012, mean ± SD in log scale, n = 3; Figure 4A). In contrast, the shoulder was absent under high-light conditions, and the slope in the same range steepened (−0.872 ± 0.011). When we focused on the single viewpoint in the green probe, the genomic distance vs contact frequency curve showed the same behavior as that of whole genome (Figure 4B).

**Figure 4.**
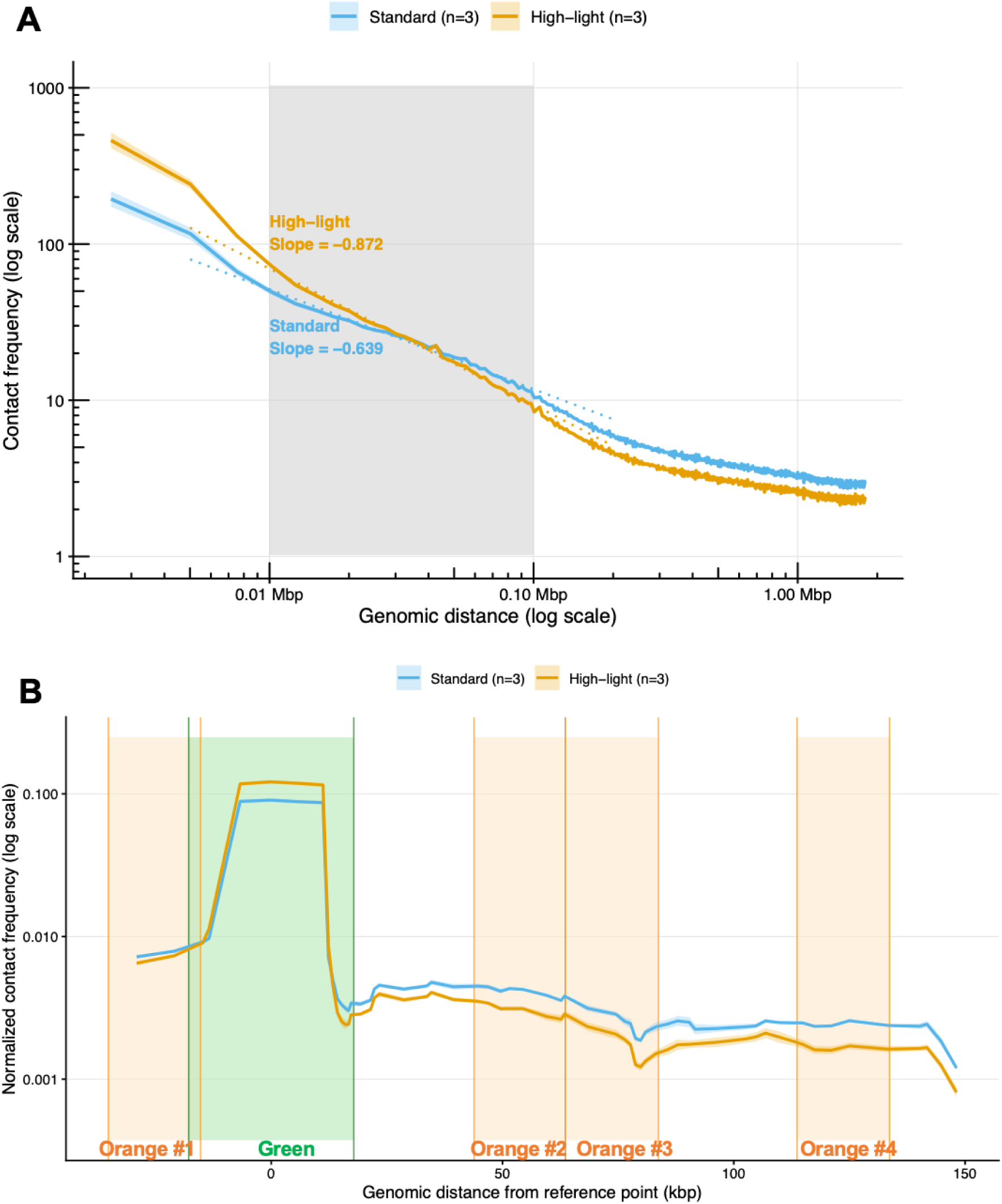
Hi-C analysis of chromosome organization under standard and high-light conditions in *Synechocystis* sp. PCC 6803. **(A)** Contact frequency as a function of genomic distance under standard and high-light conditions (150 µmol photons m⁻² s⁻¹, 80 min). Data are plotted on a log–log scale up to half the genome size (∼1.8 Mbp). Lines represent the geometric mean of three biological replicates; shading indicates ± SD. The gray shaded region highlights the 10–100 kbp range. Dotted lines indicate linear regression fits within this range, with slopes annotated. **(B)** Pseudo-4C contact frequency profile centered on the green probe reference point (genomic position 1,702,186 bp; NC_000911.1) under standard and high-light conditions (150 µmol photons m⁻² s⁻¹, 80 min). Normalized contact frequency (geometric mean of three biological replicates) is plotted against genomic distance from the reference point on a log scale; shading indicates ± SD. Colored columns indicate the genomic regions of the green (reference) and orange FISH probes (see Figure 1A for probe positions).

## DISCUSSION

### FISH-based analysis reveals local chromosome organization in *Synechocystis*

The mean number of FISH signals in the cells under standard conditions was somewhat fewer (0.4 and 1.7 copies in two biological replicates) than the qPCR-estimated GCN. This implies that overlapping of signals occurs once per cell. This can be explained by *Synechocystis* replication of only one copy of the chromosome at a time (Ohbayashi *et al*, 2019). This means that most recently replicated genomic regions detected by FISH may remain adjacent and therefore be counted as a single signal, although qPCR detects two copies. Under this assumption, FISH signal counts reflect GCN in *Synechocystis*.

The Hungarian algorithm was used to identify same-chromosome signal pairs under the assumption that, within the analyzed genomic range, green and orange FISH signals from the same chromosome are spatially clustered. The permuted dataset yielded significantly longer mean distances than the observed dataset, supporting the presence of spatial clustering and validating the use of the Hungarian algorithm (Figures 2A and 2B).

The statistically significant correlation between the genomic and spatial distance of the signal pair also supports that we preferentially measure the distances between two loci on the same DNA. Cyanobacterial nucleoid plasticity explains the low coefficient of determination (R^2^ = 0.116 by four data points and R^2^ = 0.103 by three data points).

The linear regression analysis gave a slope (β = 0.972 nm/kbp by four data points and 1.07 nm/kbp by three data points), representing the spatial distance increase per kilobase of genomic separation, which corresponds to a 300∼350-fold compaction relative to the B-form DNA contour length of 0.34 nm/bp. Notably, this scale matches the order of magnitude of the nucleoid diameter (1–2 µm) without invoking unrealistically extreme global compression.

The number of signal pairs per cell, used here as an observational proxy for GCN, showed opposing effects on the two distance metrics: a positive effect on mean distance and a negative effect on minimum distance (Supplementary Table S1 and Supplementary Figure S2). These opposing directions are consistent with a technical artifact of the pairing algorithm—as signal pair number increases, the probability of mispairing inflates mean distances, while increased signal density reduces minimum distances by chance—rather than a genuine modulation of chromosome organization by GCN. Accordingly, the interaction between GCN and genomic distance was not significant under either condition (Supplementary Table S1), indicating that the genomic–spatial distance relationship is largely independent of GCN within the range examined.

### The distribution of a single-color signal implies the partitioning of some genomic locus

The same-color nearest-neighbor distance analysis (green–green and orange–orange) assesses the spatial separation of genome copies—that is, how signals representing the same genomic locus on different chromosome copies are distributed within cells. This is conceptually and methodologically distinct from paired signal analysis (green–orange), which examines distances between different genomic loci on the same chromosome copy. To assess whether genome copies are distributed non-randomly within cells, we compared the log–log slopes of same-color nearest-neighbor distances against the theoretical expectation for complete spatial randomness (CSR) in three dimensions, which predicts a slope of −1/3 (−0.33). The log–log slopes of same-color nearest-neighbor distances shifted between conditions: slopes under standard conditions (−0.16 to −0.28) were shallower than the CSR expectation (−0.33), whereas those under high-light conditions shifted toward or beyond −0.33 (Figure 3A). Although cell-size variation precludes definitive conclusions about absolute spatial separation from CSR, the consistent directional shift between conditions supports condition-dependent changes in genome copy organization. Notably, the shallower-than-CSR slopes under standard conditions suggest that genome copies tend toward non-random spatial separation within the cell. This bears a conceptual parallel with the axial partitioning of chromosomes observed in the rod-shaped cyanobacterium *S. elongatus* (Jain *et al*, 2012; Watanabe *et al*, 2018), suggesting that spatial partitioning of genome copies may be a shared organizational feature across morphologically distinct cyanobacteria.

### Altered chromosome organization under high-light stress

Linear regression analysis revealed a substantially reduced genomic–spatial distance correlation under high-light conditions (Figure 3B). The distance-dependent decrease in effect size (r = 0.25, 0.06, and −0.05 at 53.7, 73.6, and 123.7 kbp, respectively) suggests that high-light-induced chromosome reorganization preferentially affects shorter-range spatial relationships. The altered chromosome organization under high-light is therefore unlikely to result from the slightly elevated GCN (∼7 vs. ∼6 copies per cell) and instead likely reflects a direct, high-light-induced reorganization of chromosome structure. This suggests that the preferential spatial separation of genome copies observed under standard conditions may be diminished under high-light stress, consistent with the reduced genomic–spatial distance correlation.

### Functional implications of chromosome reorganization under high-light stress

One possible explanation is that chromosome structural changes serve a protective function under high-light stress. High-light induces oxidative stress through reactive oxygen species production and direct UV damage to DNA. Altered chromosome compaction or spatial organization could potentially protect DNA from oxidative damage or facilitate access for DNA repair machinery. Alternatively, the transcriptional reprogramming required for stress responses may necessitate reorganization of the chromosome structure to modulate accessibility of stress-response genes. Further experiments combining FISH with oxidative stress markers or DNA damage sensors would help distinguish these possibilities.

Notably, in chloroplasts—organelles derived from cyanobacterial ancestors—the association of DNA with thylakoid membranes has been shown to depend on active transcription (Palomar *et al*, 2024), suggesting that transcriptional activity shapes chromosome organization in photosynthetic organelles of cyanobacterial origin. A similar principle operates in *Escherichia coli*, where coupled transcription, translation, and membrane protein insertion (transertion) anchors the nucleoid to the inner membrane, and inhibition of transcription or translation leads to rapid nucleoid detachment within minutes (Spahn *et al*, 2025). Whether analogous coupling exists between chromosome reorganization and transcriptional reprogramming in cyanobacteria under high-light stress remains an open question.

It should be noted that the present study demonstrates an association between light intensity and higher-order chromosome organization, but does not establish whether this structural change is a cause or consequence of transcriptional reprogramming or DNA protection. Determining the functional significance of the observed chromosome reorganization will require future experiments integrating chromosome conformation analysis with transcriptomic profiling and targeted perturbation of nucleoid-associated proteins.

The light-dependent chromosome reorganization observed here in *Synechocystis* shares a conceptual parallel with the chromosome reorganization reported in *S. elongatus* in response to circadian signals including compaction of the nucleoid (Smith & Williams, 2006) and rhythmic changes in DNA supercoiling (Vijayan *et al*, 2009; Woelfle *et al*, 2007), as well as reorganization under nutrient limitation and growth arrest (Dudley *et al*, 2025). Together, these observations suggest that dynamic restructuring of higher-order chromosome organization may be a common response to diverse environmental perturbations in cyanobacteria, regardless of cell morphology. Notably, because our experiments were conducted under continuous illumination, the reorganization observed here is unlikely to reflect circadian oscillation and instead points to direct light intensity as a trigger for chromosome restructuring. A recent Hi-C study in *Brucella melitensis* further reported that short-range chromosomal interactions increase upon transition from exponential to stationary phase (Huang *et al*, 2023), in contrast to the reduction observed here under high-light conditions. This difference likely reflects the distinct nature of the two stimuli—growth arrest versus acute light exposure—rather than a contradiction, and underscores that the directionality of chromosomal reorganization is shaped by the specific physiological context.

### Hi-C evidence for light-dependent chromosome reorganization

We note that FISH and Hi-C experiments were performed under different high-light intensities (300 and 150 µmol m⁻²s⁻¹, respectively). Both intensities represent substantial increases over standard conditions (40 µmol m⁻²s⁻¹). The qualitatively consistent changes observed across FISH and Hi-C analyses—including increased inter-probe distance at 53.7 kbp (FISH) and decreased short-range interactions at 10–100 kbp (Hi-C and pseudo-capture Hi-C)—suggest that chromosome reorganization is a general response to elevated light intensity rather than a threshold effect specific to a particular intensity.

### Limitations and future directions

Although our FISH-based approach successfully revealed local chromosome organization in *Synechocystis* at single-cell resolution, two technical limitations should be acknowledged. First, our analysis was limited to four genomic distances (25–124 kbp) within a single chromosomal region, precluding assessment of shorter- or longer-range organization. Second, assigning signal pairs in multi-copy genomes carries inherent uncertainty; while permutation testing and genomic distance correlation analysis provide validation, we cannot definitively confirm that all detected pairs originate from the same chromosome copy.

These limitations could be addressed by two complementary strategies. Multicolor FISH with three or more spectrally distinct probes would enable same-chromosome identification by color combination, eliminating pairing ambiguity and extending the range of analyzable genomic distances. Alternatively, two-color live-cell imaging would provide dynamic information inaccessible to fixed-cell FISH: loci moving in concert are likely to reside on the same chromosome copy, offering an orthogonal approach to pairing validation. Since bacterial nucleoids are dynamically shaped by cellular activities such as transcription and translation (Spahn *et al*, 2025; Kuzminov, 2024; Papagiannakis *et al*, 2025), live imaging of cyanobacterial nucleoids will ultimately be essential for understanding the temporal dimension of chromosome organization revealed here by FISH.

## MATERIALS AND METHODS

### Experimental Design

To investigate whether higher-order chromosome organization changes in response to light intensity in *Synechocystis* sp. PCC 6803, we employed an integrative approach combining two-color FISH and Hi-C analysis. FISH experiments were performed under standard light conditions (50 µmol photons m⁻² s⁻¹) and high-light conditions (300 µmol photons m⁻² s⁻¹) using probe pairs targeting genomic loci separated by 25.3–123.7 kbp. Each FISH experiment was independently replicated twice using biological replicates. Hi-C analysis was performed under standard (50 µmol photons m⁻² s⁻¹) and high-light (150 µmol photons m⁻² s⁻¹) conditions to provide population-level contact frequency data complementary to the single-cell FISH measurements. Although the two methods were performed under different high-light intensities, the convergent findings across these conditions strengthen the conclusion that the observed chromosome reorganization reflects a directional response to elevated light intensity.

### Bacterial strains and culture conditions

For the standard conditions, the glucose-tolerant strain of *Synechocystis* sp. PCC 6803 (Williams, 1988) was cultivated in BG-11_0_ liquid medium (Rippka *et al*, 1979) containing 5 mM NH_4_Cl and buffered with 20 mM HEPES-KOH (pH 7.8) under continuous exposure to white light (40 µmol m^−2^s^−1^) and bubbled with air containing 1% CO_2_. Cells in the linear growing phase (OD730 ≈ 2) were harvested for fixation (standard conditions sample) or exposed to stronger white light (300 µmol m^-2^s^-1^) for 80 minutes and harvested (high-light-condition sample). For high-light conditions of Hi-C experiments, cells were exposed to 150 µmol m⁻²s⁻¹ for 80 minutes and harvested. For the phosphate-depleted condition, cells with OD_730_ ≈ 0.6 were collected and washed twice with phosphate-depleted BG-11 (BG-11_0_ liquid medium without phosphate containing 5 mM NH_4_Cl and buffered with 20 mM HEPES-KOH (pH 7.8)), then diluted to OD_730_ = 0.1 by phosphate-depleted BG-11 and cultivated for six days.

### Cell fixation for FISH

Two biological replicates were sampled for all conditions. Ten milliliters of cultures of *Synechocystis* sp. PCC 6803 in the conditions mentioned above was mixed with 30 mL of freshly made 4% Paraformaldehyde (Merck Millipore, Darmstadt, Germany) in PBS, and the mixture was shaken for an hour at room temperature. Then cells were collected by centrifugation at 2264 ×g for 2 min and washed with PBS twice. Finally, the cells were suspended in EtOH: PBS = 1:1(w/w) solution and stored at −30°C until use. For each replicate, the fixed cell suspension was divided into aliquots and independently hybridized with each probe combination as described in Results.

### Ploidy determination

Genomic DNA was extracted from fixed cells as follows: Fixed cells were suspended in 100 µL of TE buffer (10 mM Tris-HCl pH 7.5 and 1 mM EDTA), and the cell number was counted with Thoma’s Counting Chamber. Cells were lysed by mixing 45 µL of cell suspension, 5 µL of 10% Sodium Dodecyl Sulfate (SDS), and 100 µL of glass beads (G1145, Sigma-Aldrich, St. Louis, Missouri), followed by shaking for 3 minutes with a twin mixer (TM-282, AS ONE Co., Osaka, Japan). This method disrupted >99% of cells as confirmed by microscopy. 100 ng of λstyI DNA marker (2.5 µL of 40 ng/µL stock, Nippon Gene) was added as an internal standard for the recovery rate calculation. The lysate was diluted with TE buffer, and an aliquot was transferred to a new tube to avoid glass beads. The lysate was treated with RNase A (Nippon Gene) at 37°C for 1 h, followed by SDS (to a final concentration of 1%) and proteinase K (to a final concentration of 50 µg/mL, Kanto Chemical Co., Tokyo, Japan) at 65°C for 6 h for protein digestion and decrosslinking. DNA was purified by phenol-chloroform extraction and ethanol precipitation, and dissolved in 25 µL of TE buffer.

Absolute quantification of genomic DNA was performed according to the method described by (Griese *et al*, 2011) with modifications. Briefly, the molecular number of genomic DNA was quantified by qPCR using 1 kbp standard DNA fragments, and the molecular number of λDNA before (100 ng input) and after purification was quantified.

The original molecular number of genomic DNA in the cell lysate was back-calculated using this recovery rate. Finally, ploidy (GCN per cell) was determined by dividing the total molecular number of genomic DNA by the cell number.

### Cell permeabilization for in situ hybridization

The fixed cells were washed and suspended with GTE buffer (50 mM glucose, 20 mM Tris HCl pH7.5, and 10 mM EDTA). 150 µL of the cell suspension was dropped onto a MAS-coated slide glass (Matsunami glass, Tokyo, Japan). After 1 hour of incubation at room temperature, slide glasses were soaked with GTE buffer at 37°C. The cell wall was then digested by incubation at 37°C for 30 minutes in GTE with 0.2% 2-mercaptoethanol and 50 µg/mL of hen-egg lysozyme (FUJIFILM Wako Pure Chemical Corp., Osaka, Japan). After digestion, the slides were soaked in 5 mg/mL CuSO_4_·5H_2_O solution at room temperature for 30 minutes to quench the autofluorescence. They were then sequentially soaked in

70%, 85%, and 100% ethanol, at room temperature and air-dried. For the denaturation, the slides were soaked in 50% formamide / 2×SSC at 72°C for 3 minutes, then immediately transferred into ice-chilled 70%, 85%, and 100% ethanol for 3 minutes each. Finally, the slides were air-dried and waited for the hybridization step.

### Fluorescent labeling of the FISH probe

Template of fluorescent probes were either CS0508, the cosmid of *Synechocystis* sp. PCC 6803 genome (Kotani *et al*, 1994) or PCR products amplified with PrimeSTAR® LongSeq DNA Polymerase (Takara Bio, Shiga, Japan) and primer oligonucleotides listed in Table S2. Fluorescent probes were synthesized using a nick translation kit (Roche, Basel, Switzerland) following the manufacturer’s instructions. dUTP conjugated with Spectrum™ Green (Abbott, Chicago, Illinois) was used for CS0508, and Spectrum™ Orange (Abbott) was used for the PCR products.

## FISH

Hybridization and washing were conducted as described in (Kawamura *et al*, 2012) with two modifications: a labeled probe equivalent to 10 ng of template DNA and 5 µg of salmon sperm DNA per slide were mixed and ethanol-precipitated. Slide and probe denaturation were conducted separately.

### Microscopic analysis

Fluorescent images were acquired using a laser confocal microscope equipped with Airyscan 2 (LSM900, Zeiss) and a 63× oil immersion objective (NA 1.4). DAPI, Spectrum Green, and Spectrum Orange were excited at 405 nm, 488 nm, and 561 nm, respectively, and detected at 400–505 nm, 450–550 nm, and 560–600 nm, respectively. Three fluorescent signals were imaged sequentially. Z-stacks of approximately 20 slices with 0.1 µm intervals were obtained in Airyscan mode. Each section was scanned 8 times and averaged to improve the signal-to-noise ratio. The raw data were processed using ZEN. Images were acquired with a pixel size of 0.0165 µm × 0.0165 µm in the lateral (xy) plane and a z-step size of 0.1 µm along the optical axis. Signal analysis was performed based on previous studies (Kuroda *et al*, 2004; Tanabe *et al*, 2002) with some modifications. Image files in the CZI format were analyzed with FIJI (Schindelin *et al*, 2012). Fluorescent peak centers (x, y, z in pixels) of Spectrum™ Green and Spectrum™ Orange were determined as follows: To minimize measurement bias, images were split by one cell and quantified under blinded conditions, with information on the experimental condition, probe identity, and replicate number concealed. The order of quantification was also randomized.

Fluorescent peak centers were determined using a semi-automated approach with a FIJI macro: The operator specified the region around the visible signal, and the macro searched for the local intensity maximum within a cylindrical region of 7-pixel radius and ±2 slices in z centered on the specified position by applying the Find Maxima function to each z section. Note that the Spectrum Green and Spectrum Orange signals within the same cell were detected in the same session, which may have introduced a degree of observer bias in peak localization. The 3D Euclidean distances between the two signals were measured.

### Statistical analysis and algorithm for pairing FISH signals

Statistical analysis of FISH signals was performed using R version 4.5.2. To pair Spectrum™ Green and Spectrum™ Orange signals within each cell, the Hungarian algorithm was employed to identify the pairing configuration that minimized the total sum of inter-probe distances across all pairs in the cell. The algorithm was implemented using the “clue” package version 0.3-66 (Hornik, 2005). When the number of green and orange signals was unequal, the number of pairs was limited to the smaller of the two. For the permutation test, all possible reassignments of color labels (green vs. orange) to the observed spatial coordinates within each cell were generated. For each permutation, pairs were identified using the Hungarian algorithm as described above, and the mean inter-probe distance was calculated.

A simple linear regression model was used to estimate the mean signal distances per cell as a function of the genomic distances of the probe pair. To examine whether the genomic–spatial distance relationship was modulated by genome copy number (GCN), multiple regression models with an interaction term were fitted using both mean and minimum Green–Orange distances as response variables. The predictors were genomic distance and the number of signal pairs per cell (a proxy for GCN), both mean-centered prior to model fitting to improve interpretability of the main effects. Models were fitted separately for standard and high-light conditions. The interaction term (genomic distance × number of signal pairs) was used to test whether the slope of the genomic–spatial distance relationship varied with GCN. Full model results are provided in Supplementary Table S1.

To compare mean inter-probe distances between observed data and color-label permutation controls, a one-sided paired Wilcoxon signed-rank test (alternative hypothesis observed distances are shorter than permutation controls) using the wilcox_test() function (paired = TRUE, alternative = ’less’) in the stats package was performed. To compare spatial distances between standard and high-light conditions at each genomic separation, Mann–Whitney tests (Wilcoxon rank sum tests) were performed using the wilcox_test() function. Significance thresholds for multiple comparisons across three genomic separations (53.7, 73.6, and 123.7 kbp) were adjusted using the Holm procedure. Effect sizes were quantified as rank-biserial correlation coefficients using the wilcox_effsize() function in the rstatix package (Kassambara, 2020).

### AI-assisted Scripts for statistics and data formatting

Statistical scripts and data formatting were assisted using ChatGPT (GPT-5.2; OpenAI) and Claude (Claude Sonnet 4.6; Anthropic, claude.ai). All AI-assisted content was reviewed and approved by the authors. No primary data interpretation or hypothesis generation was performed by the model.

### Hi-C experiment and analysis

Fixation and cell lysis of *Synechocystis* sp. PCC 6803 for Hi-C was performed as described in a previous study (Kariyazono & Osanai, 2024). Hi-C library preparation was performed according to Takemata and Bell (Takemata & Bell, 2021), using HindIII for restriction digestion. Sequencing (150 bp paired-end) was performed on a HiSeq X (Illumina) by an external sequencing service.

Raw sequencing reads were processed with fastp v0.23.4 (Chen *et al*, 2018) with default parameters for quality control and adapter trimming. Trimmed reads were mapped to the *Synechocystis* sp. PCC 6803 chromosome (NC_000911.1) using BWA-MEM (Li & Durbin, 2009) via hicBuildMatrix (HiCExplorer v3.7.5 (Wolff *et al*, 2020)) with a bin size of 2,511 bp. Bias correction was applied using hicCorrectMatrix with Knight-Ruiz (KR) normalization. Contact frequency as a function of genomic distance was computed using hicPlotDistVsCounts. Pseudo-4C analysis was performed using chicViewpointBackgroundModel and chicViewpoint with the green probe region (NC_000911.1: 1,684,343–1,720,028) as the reference point, followed by data export using chicExportData.

### Data, code, and materials availability: Data availability and Accession Numbers

Raw sequencing reads and processed data (KR-corrected contact matrices and distance vs. contact frequency data) were deposited in the Gene Expression Omnibus (GEO; accession number GSE326282) and the Sequence Read Archive (SRA; accession number PRJNA1438009). Raw microscopy images and all custom R scripts used for FISH signal analysis and statistical analyses are available from the corresponding author upon reasonable request. This study did not generate any new materials.

## Author contributions

Conceptualization: R.K.,T.O.; Methodology: R.K.,H.T.; Investigation: R.K.,H.T.; Visualization: R.K.; Supervision: T.O.; Writing—original draft: R.K.; Writing—review & editing: R.K.,H.T.,T.O.

## Competing interests

All other authors declare they have no competing interests.

## Acknowledgments

We thank Prof. Shusei Sato for kindly giving the cosmid CS0508 and Prof. Michiyo Honda for kindly providing access to the microscope for the optimization of the FISH experiment. We also thank Dr. Mieko Kono for providing the detailed method for bacterial FISH, and Dr Yohei Terai and Shiho Takahashi-Kariyazono for kindly providing equipment for NGS library preparation. Hi-C data processing were performed on the NIG supercomputer at ROIS National Institute of Genetics. The authors would like to thank Enago (www.enago.jp) for the English language review.

## Funding

Grant-in-Aid for JSPS Fellows grant JP24KJ2047 (K.R.)

JST-GteX of the Japan Science and Technology Agency JPMJGX23B0 (T.O.)

## Notes

### Competing Interest Statement

The authors have declared no competing interest.

### Summary of Updates

Abstract, introduction, and discussion have been revised.

