## Supplemental FIgures S1 to S4 and Supplemental Tables S1 and S2 for "Light-dependent changes in the higher-order DNA structure of the cyanobacterium *Synechocystis* sp. PCC6803"

[1]School of Agriculture, Meiji University, Kawasaki, Japan [2]Research Center for Integrative Evolutionary Science, SOKENDAI, Hayama, Japan.
*Ryo Kariyazono and Takashi Osanai

**This PDF file includes:**

Figures S1 to S4
Tables S1 to S2

**Figures**

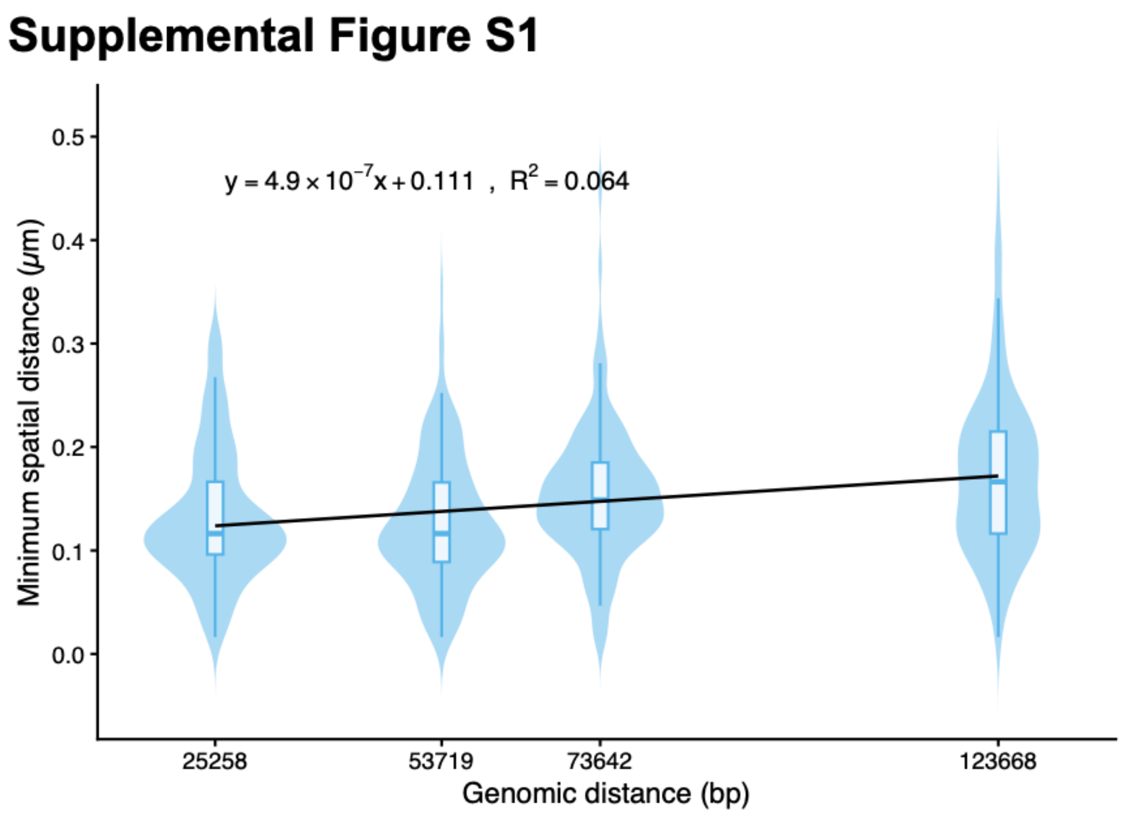


**Figure S1.** Minimum spatial distance between paired FISH signals as a function of genomic distance.

The minimum Green–Orange spatial distance per cell (minimum distance) is plotted against the genomic distance of the probe pair for standard (blue) and high-light (orange) conditions. Data from both biological replicates are pooled. Distributions are shown as violin plots with overlaid box plots (median and interquartile range). Linear regression lines for each condition are shown. p-values and rank-biserial correlation coefficients (r) from Mann–Whitney tests comparing standard and high-light conditions at each genomic separation are indicated above each distribution.

#### Supplemental Figure S2

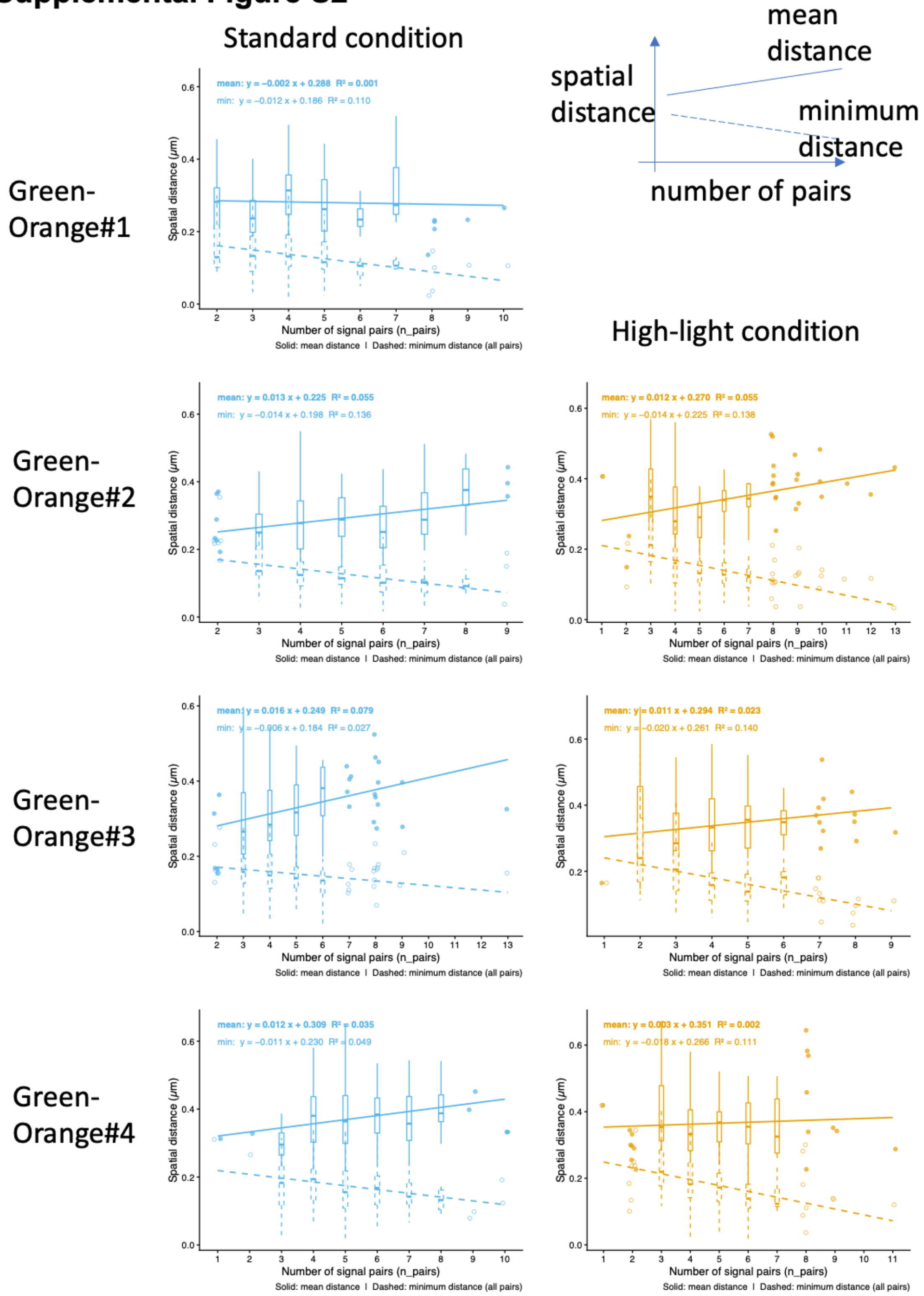


**Figure S2.** Effect of signal pair number on mean and minimum spatial distances. Mean spatial distance (solid line) and minimum spatial distance (dashed line) per cell are plotted against the number of Green–Orange signal pairs per cell, shown separately for each probe under standard (blue) and high-light (orange) conditions. Groups with ≥10 cells are shown as box plots; groups with <10 cells are shown as jittered points. Linear regression lines and fit statistics (slope, intercept, $R^2$) are shown for each metric. Note the opposing directions of the regression slopes for mean and minimum distances, consistent with a technical artifact of the pairing algorithm rather than a genuine biological effect of genome copy number.

### Supplemental Figure S3

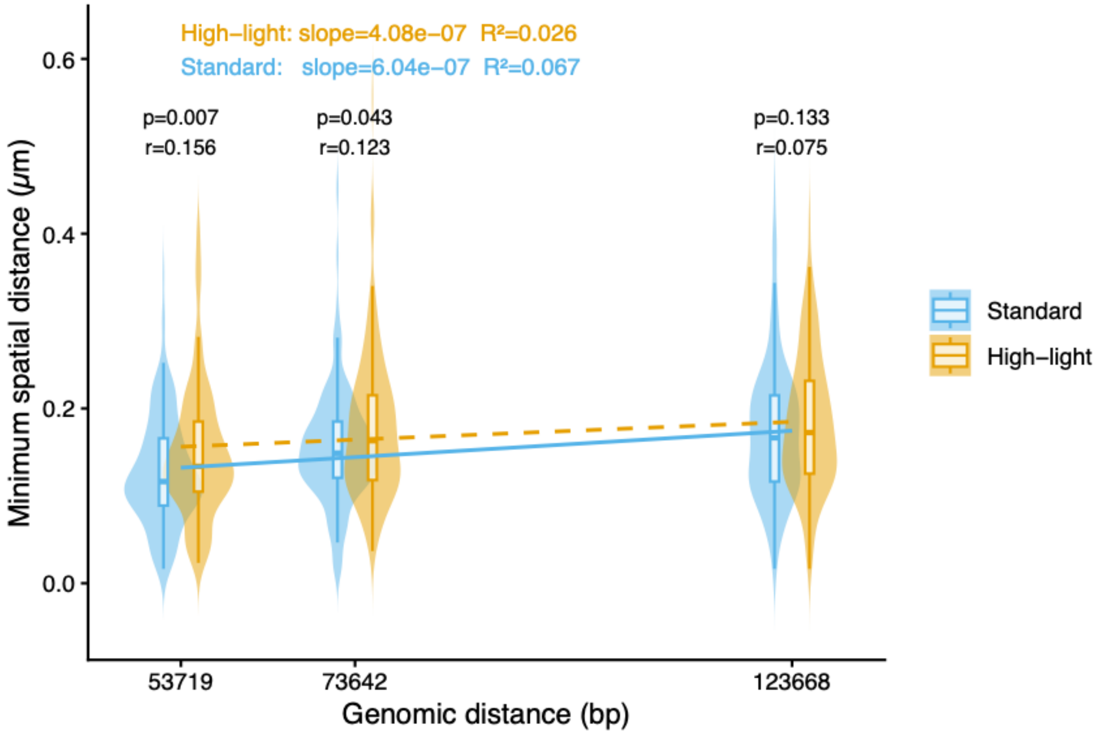


**Figure S3.** Minimum spatial distance between paired FISH signals as a function of genomic distance under standard and high-light conditions. Layout as in Figure S1, except that standard (blue) and high-light (orange) conditions are compared across three probe pairs (53.7, 73.6, and 123.7 kbp). p-values and rank-biserial correlation coefficients (r) from Mann–Whitney tests comparing the two conditions at each genomic separation are indicated above each distribution.

#### Supplemental Figure S4

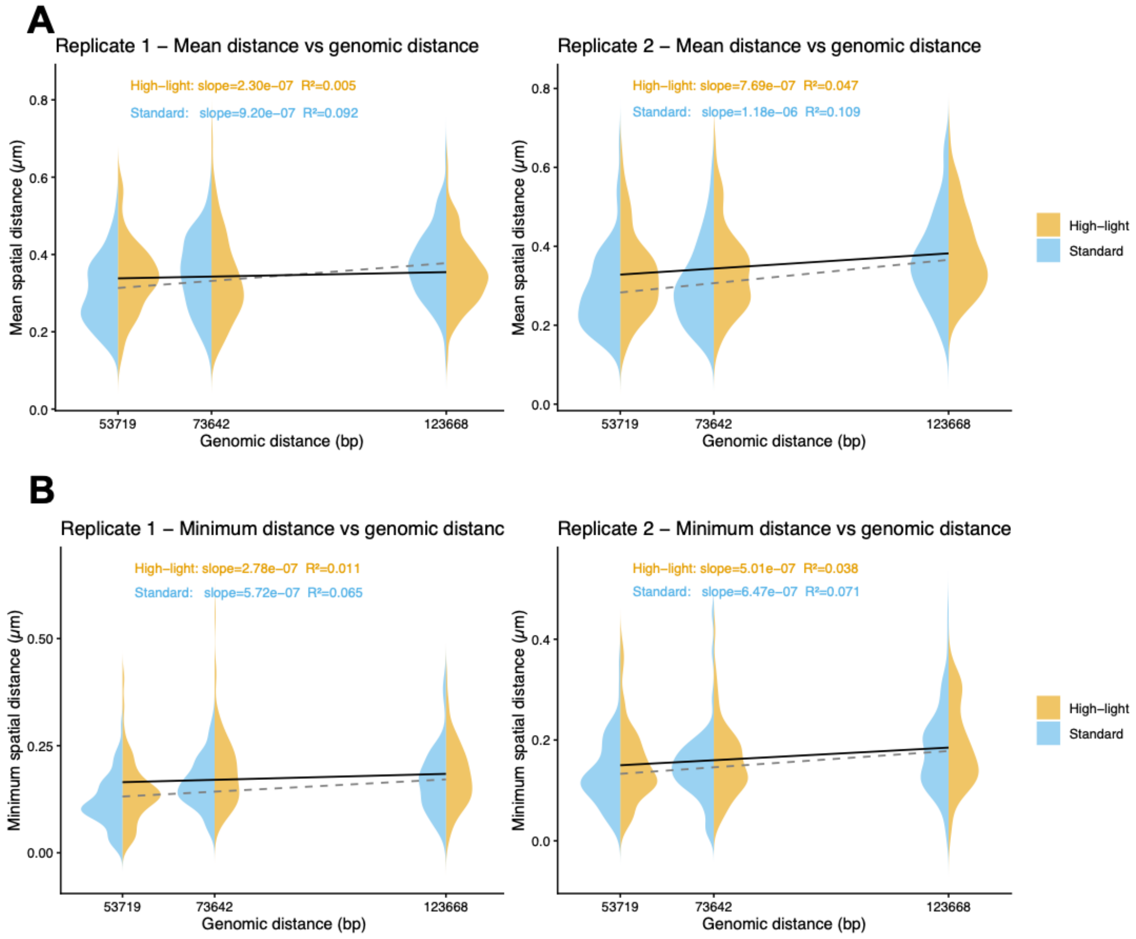


**Figure S4.** Replicate-specific spatial distance analyses. (A) Mean spatial distance between paired Green–Orange FISH signals as a function of genomic distance, shown separately for each biological replicate (rep 1 and rep 2). Standard (blue) and high-light (orange) conditions are displayed as split violin plots. Linear regression fit statistics (slope, $R^2$) are shown for each condition. (B) Minimum spatial distance as a function of genomic distance, shown separately for each biological replicate. Layout as in (A).

**Tables**

**Table S1** Interaction model results for spatial distance as a function of genomic distance and genome copy number.

| | | Standard | | High-light | |
|---|---|---|---|---|---|
| metrics | **Term** | **Standard β (SE)** | **Standard p** | **High-light β (SE)** | **High-light p** |
| (a)Mean distance | **Intercept** | 3.326e-01 (4.435e-03) | <0.001 | 3.486e-01 (4.400e-03) | <0.001 |
| | **genomic_dist** | 1.049e-06 (1.449e-07) | <0.001 | 4.905e-07 (1.423e-07) | <0.001 |
| | **n_pairs** | 1.335e-02 (2.731e-03) | <0.001 | 7.384e-03 (2.673e-03) | 0.00595 |
| | **genomic_dist × n_pairs** | -2.765e-08 (9.102e-08) | 0.761 | -1.229e-07 (8.649e-08) | 0.156 |
| | **$R^2$** | 0.149 | | 0.042 | |
| | **n** | 463 | | 516 | |
| (b)minimum distance | **Intercept** | 1.544e-01 (3.141e-03) | <0.001 | 1.681e-01 (3.210e-03) | <0.001 |
| | **genomic_dist** | 6.222e-07 (1.026e-07) | <0.001 | 3.281e-07 (1.038e-07) | 0.00167 |
| | **n_pairs** | -1.070e-02 (1.934e-03) | <0.001 | -1.709e-02 (1.950e-03) | <0.001 |
| | **genomic_dist × n_pairs** | 1.443e-08 (6.446e-08) | 0.823 | -3.696e-08 (6.309e-08) | 0.558 |
| | **$R^2$** | 0.127 | | 0.156 | |
| | **n** | 463 | | 516 | |

.

**Table S2.** Oligonucleotides used in this study

| Oligo ID | sequence 5'->3' | Purpose | source |
|---|---|---|---|
| Standard 1 | CGCGGATTACCCTACCAG ACCGGCGATC | determination of ploidy by qPCR (amplificaton of Standard DNA fragment) | K Zerulla et al 2016 |
| Standard 1 | GGTCCCAATACGGTTGGT AAGCCCTTCG | determination of ploidy by qPCR (amplificaton of Standard DNA fragment) | K Zerulla et al 2016 |
| Analysis 1 | GTCCTATCTAATCGCTGT GGTAGCCAACCGC | determination of ploidy by qPCR (qPCR primer) | K Zerulla et al 2016 |
| Analysis 1 | CTGTCCCCACTGCCAAAG CTTAAGCCC | determination of ploidy by qPCR (qPCR primer) | K Zerulla et al 2016 |
| Standard 2 | CCCCACCTCTCCCCGTCG TCCCC | determination of ploidy by qPCR (amplificaton of Standard DNA fragment) | K Zerulla et al 2016 |
| Standard 2 | GCTGCTTCCTGCATAGCC ACGGGGG | determination of ploidy by qPCR (amplificaton of Standard DNA fragment) | K Zerulla et al 2016 |
| Analysis 2 | GGACCATGCCCCTATTGC CTACGACTGGC | determination of ploidy by qPCR (qPCR primer) | K Zerulla et al 2016 |
| Analysis 2 | CGATCGCCCCTGCCAATT CCACCC | determination of ploidy by qPCR (qPCR primer) | K Zerulla et al 2016 |
| Standard 3 | GGACCCGCCTCTATCCTT TAACCAAACTCAGAGCC | determination of ploidy by qPCR (amplificaton of Standard DNA fragment) | K Zerulla et al 2016 |
| Standard 3 | GTAGAAATCATTGCCCAT CACCAAAGTATCCTCAAT GG | determination of ploidy by qPCR (amplificaton of Standard DNA fragment) | K Zerulla et al 2016 |
| Analysis 3 | CCAACAAACCAAAGATAA TCCTGATTGGTTTCAGGG | determination of ploidy by qPCR (qPCR primer) | K Zerulla et al 2016 |
| Analysis 3 | CAGAAAAGTCAGTAATTCT GCCCTGGGCGTCG | determination of ploidy by qPCR (qPCR primer) | K Zerulla et al 2016 |
| qPCRlambda_styI_19k_F | GATACGGCGTGAACGTCT TC | determination of ploidy by qPCR (qPCR primer for lambda DNA) | in this study |
| qPCRlambda_styI_19k_R | CATGCTTCTCCAGTGCAT CC | determination of ploidy by qPCR (qPCR primer for lambda DNA) | in this study |
| FISH_hoxregion_F | TAGGATCCACGGCCCTAG CCAAAAGAGTGGTGTGG | Long PCR primer for template of FISH Orange#1 | in this study |
| FISH_hoxregion_R | TTACCCAGGTCATCGGGC CTGTAATTGACGCCCAG | Long PCR primer for template of FISH Orange#1 | in this study |
| FISH_1766k_1F | GGGCGATCGCCAATTTCC CCAACGGCTGCAAATCC | Long PCR primer for template of FISH Orange#2 | in this study |
| FISH_1766k_1R | CGGGCATTGCCTGTTATC CCAACCATGGTGATAACG TGG | Long PCR primer for template of FISH Orange#2 | in this study |

| | | | |
|---|---|---|---|
| FISH_1766 k_2F | ATGCCCGCATAGCCCTGG AGGTAAAGGGGATTATTA GC | Long PCR primer for template of FISH Orange#3 | in this study |
| FISH_1766 k_2R | GAGCCTGGCACCCGTTTC TCTGCCGGTATCCAGTG | Long PCR primer for template of FISH Orange#3 | in this study |
| FISH_1816 _2F | GCGATGTGGAAGGATTGC GGGGTTTCATTCCCCGAT C | Long PCR primer for template of FISH Orange#4 | in this study |
| FISH_1816 _2R | ACCACTAACTTTCCCGAG TACGTTGAACCGCCACCC | Long PCR primer for template of FISH Orange#4 | in this study |
